# Genetic diversity of *Bemisia tabaci* Genn. characterized by analysis of ISSR and cytochrome c oxidase subunit I at Qassim, Saudia Arabia

**DOI:** 10.1101/2020.12.29.424654

**Authors:** Nagdy F. Abdel-Baky, J. K. Brown, M. A. Aldeghairi, M. I. Motawei, Medhat Rehan

**Author notes:** Corresponding Author: **Mailing address:** Department of Plant Production and Protection, College of Agriculture and Veterinary Medicine, Qassim University, Saudi Arabia. **E-mail:**. Or. **E-mail:** Or.

## Abstract

Problems of Bemisia tabaci (Gennadius) (Hemiptera: Aleyrodidae) that increased and escalated in the last 40 years seem to be related to one or more aggressive biotypes that appeared to spread steadily worldwide. As well, some biological characteristics of B. tabaci have led some entomologists to change and multiply their methodology to update with the change in the pest genetic structures. This study is the 1st of its kind in Qassim region in KSA in respect of *B. tabaci* biotypes. Four identification methods (Squash Silverleaf Symptoms (SSL), cross mating, Cytochrome oxidase subunit I (COI) sequences, and ISSR-PCR analysis) were carried out to determine the biotypes of B. tabaci at Qassim regions. Slight SSL symptoms were observed with varying degrees on squash leaves caused by *B. tabaci* population at Qassim, KSA. Cross-mating among the populations that have the same or similar genetic structures produced fertilized offspring, females and males with higher sex ratios in favor of females, and produced a higher number of eggs. Whereas, B. tabaci populations that varied greatly in their genetic structures produced unfertilized eggs, which produce males only. In the same trend, ISSR-PCR analysis revealed that B. tabaci populations at Qassim regions varied genetically and gathered into four genetic groups. In conclusion, COI analysis is a perfect tool for classification between biotypes in B. tabaci. Therefore, this study declares that *B. tabaci* that colonized and infest Qassim horticulture has not the same genetic structures but belonging to B biotypes. It could be named as *Bemisia* species complex.

## Introduction

*Bemisia tabaci* (Gennadius) is considered one of the most key and important whiteflies in the world which originated from warm to hot climates. *B. tabaci* is classified under family Aleyrodidae (Sternorryncha: Hemiptera) (1)(2). Of the nearly 1450 described whitefly species (3), *B. tabaci* is the most widely distributed, and consider the greatest importance as a harmful insect pest in the tropics, subtropics, and the Mediterranean Basin. Globally, *B. tabaci* is a cosmopolitan pest of an extended host range of economic plants that colonizes over 900 perennial and annual plants, causing direct damage due to feeding and indirect damage by transmitting a great number of plant viruses (4)(5)(6) (7).

The classical taxonomic relationships of *B. tabaci* species complex, which have been confused for many years, remain frustrating (8). Because of the difficulty in preparing a good adult slide mounts, Scientists differentiated between whiteflies and Bemisia species based on pupal morphological characters (9)(10)(3). Despite the identification of Bemisia species based on the 4th pupal instar, another difficulty may be faced because of variations in the setae or shape of the pupal case (8). The pupal setae, size, shape, and other morphological characters are affected by the plant host and environmental factors (3). Additionally, since the pupa morphological characteristics exhibit numerous variations, they are not reliable to distinguish between *B. tabaci* biotypes (11). Therefore, accurate identification of *B. tabaci* species complex is a prerequisite for their effective control methods (12). Bemisia species are rich in their biological variants and lack external distinguishing morphological characteristics (13)(8).

*Bemisia tabaci* is a species complex (14) that includes about 24 different biotypes (15). These biotypes vary in certain biological characters *i*.*e*., dispersal, reproductive rate, host range, insecticide resistance; virus vectorially, and certain plant physiological disorders (16)(8)(17)(18). High genetic polymorphism has been confirmed in the whitefly *B. tabaci* populations based on several types of molecular analyses (15). Despite significant genetic variation among its biotypes and haplotypes are morphologically indistinguishable, lacking clear-cut morphological characters in either the pupae or the adults (19)(3). Genetic polymorphisms were first demonstrated based on unique esterase banding patterns (20), when applied, yielded numerous haplotypes (21)(22)(12). This nomenclature is still in use by the scientific community to refer to particular biotypes and for historical reasons.

To distinguish between *B. tabaci* biotypes, (23) concluded at least six identification methods namely; 1) plant physiological disorders, 2) biological characteristics, 3) crossing experiments and mating behavior, 4) morphological characters, 5) morphometrics characters, and 6) DNA molecular analysis, in addition to the shape and number of the bacterial-like organisms that live in Bemisia. The authors emphasized that at least two of these methods must be combined in Bemisia biotypes identification, particularly, genetic, physiological, and morphological examination (24)(19)(11).

Regarding *B. tabaci* biotypes, which harbored plants in Saudi Arabia, there are no significant efforts were carried out to verify these biotypes. To our best knowledge, this study considers the first studies regarding the determination of *B. tabaci* biotypes in KSA, particularly in the Qassim region.

Therefore, our research objectives in this study were: i) to determine the sweet potato whitefly (*Bemisia tabaci*) biotypes that spread in Qassim region; ii) to determine the host plants associated with these biotypes; iii) to identify the most important and harmful biotype through determining the population density; iv) to use DNA markers to differentiate between the biotypes that closely related and v) to link between DNA fingerprinting analysis and the ability of biotype to induce certain plant physiological disorders

## Materials and Methods

### Survey and Systematic Sampling of *B. tabaci*

As known, *B. tabaci* invades plants either in open fields or in the greenhouses. Most of *B. tabaci* samples were taken from plants grown under greenhouses since farmers mainly cultivate their vegetable crops under greenhouse according to weather condition.

Eight vegetable crops namely; lettuce (*Lactuca sativa* L.; Asteraceae), cabbage (*Brassica oleracea* L.; Brassicaceae or Cruciferae), cauliflower (*Brassica oleracea var. botrytis*; Brassicaceae or Cruciferae), eggplant (*Solanum melongena* L.; Solanaceae), pepper (*Capsicum annuum* L.; Solanaceae), potato (*Solanum tuberosum* L.; Solanaceae), tomato (*Solanum lycopersicum* L.; Solanaceae) and cucumber (*Cucumis sativus* L.; Cucurbitaceae) were chosen from 10 greenhouses located in different locations in Qassim region, KSA (Table 1) during the year 2010. *B. tabaci* adults were collected in the early morning by two collecting apparatus namely aspirators and electronic suction tarps. Samples were divided into two parts, one part was left live to establish new lab colonies to complete other studies, while the others were preserved in 100% ethanol and stored at −20°C until DNA extraction. At least, 100 *B. tabaci* individuals were present in each collection, otherwise, one-fourth of selected individuals from each collection were randomly selected for the biotype determination.

**Table 1.** Location and Host Plants for the sweet potato whitefly, *B. tabaci*, which collected from the Al-Qassim region.

| Code | Location | Host plant | Plant Scientific names |
| --- | --- | --- | --- |
| V1 | Agri. Exp. St., Qassim Univ. | Lettuce | *Lactuca sativa* L<br>Asteraceae |
| V4 | Agri. Exp. St., Qassim Univ. | Cucumber | *Cucumis sativus* L.<br>Cucurbitaceae |
| V7 | Ouen Al-Gawa | Cabbage | *Brassica oleracea* L.;<br>Brassicaceae or Cruciferae |
| V6 | Ouen Al-Gawa | Potato | *Solanum tuberosum* L.;<br>Solanaceae |
| V10 | Al-Melada | Tomato | *Lycopersicon esculentum* L |
| V9 | Unayzah | Eggplant | *Solanum melongena* L.;<br>Solanaceae |
| V2 | Al-Bukariyah | Cabbage | *Brassica oleracea* L.;<br>Brassicaceae or Cruciferae |
| V3 | Al-Bukariyah | Cauliflower | *Brassica oleracea* var.<br>*botrytis*; Brassicaceae or<br>Cruciferae |
| V5 | Al-Badai | Sweet pepper | *Capsicum annuum* L.;<br>Solanaceae |
| V8 | Al-Midhnab | Cabbage | *Brassica oleracea* L.;<br>Brassicaceae or Cruciferae |

### Establishing colonies in *B. tabaci* collected populations

Live individuals of *B. tabaci* adults that previously collected, were used to establish a colony for each population under lab condition. Adults of each population were introduced into large screen cages (40 × 40 × 30 cm) on white kidney beans (*Phaseolus vulgaris* L.) placed in small plastic pots. Both insects and plants were maintained at 26 ± 2 °C, 65% RH with a 14:10 (L: D) photoperiod. To produce plants with a heavy infestation from *B. tabaci* immature, newly emerged plants were infested with *B. tabaci* adults for 48 h to deposit their eggs, resulting in at least 50–100 eggs/leaf. Insects of each population were left to produce a huge number to achieve the following trials.

### Population density of *B. tabaci* associated with plant host

The population density of *B. tabaci* associated with each plant host was estimated. Twenty-five leaves from each plant host/greenhouse were chosen in random assimilated all leaves on the main plant stem (from top to bottom). Chosen leaves were cut off, inserted in plastic bags, and transformed into the lab for investigation. The number of *B. tabaci* eggs and nymphs/cm2/leaf/plant was counted under binocular by dividing the plant leaf into sectors, each sector equal 1cm^2^. The same procedure was repeated with each plant host/greenhouse for each location.

### Plant physiological disorder (Squash Silverleaf (SSL) bioassay)

To study a physiological disorder disease associated with feeding activities of certain *B. tabaci* biotypes, squash plants were seeded in black plastic pots under laboratory conditions. About 55 pots were prepared for this purpose. After the emergence of the first three true leaves, 10 pairs of each *B. tabaci* population were introduced for 48 hours until getting insect eggs. Infested squash plants for each population were observed daily for at least the emergence of F1 adults and the symptoms of squash silvering leaf (SSL) were also observed. Each population was replicated 5 times, as well as, five pots as a control (free of insects).

### Cross mating trials between collected *B. tabaci* populations

Ten individuals of newly emerged *B. tabaci* were separated from each population that reared at lab conditions and recognized to males and females. One male and female (of each population) were caged on kidney bean plants under a clip cage, and let for mating. The trial for each population was replicated 5 times. After 3 days, insect parents were removed, and laid eggs were counted, observed until the emergence of the adult. Immediately after the adults were removed from the plants, all leaves of the plants were examined using a hand lens to count the eggs laid. The plants were then cultured until all the eggs had either completed development to adults or died, and the resultant adults in each replicate were counted and sexed. From these recordings, the number of the deposited eggs, and sex ratios of viable offspring were collected for each replicate. It means that progeny and sex ratio of F1 were taken into account for each population.

### DNA extractions and PCR reactions (ISSR assay)

DNA from *B. tabaci* insects was extracted (25). The Inter Simple Sequence Repeat (ISSR) PCR method was carried out (26). Amplification was carried out in 25 μl reaction volumes, containing 1X Taq polymerase buffer (50 mM KCl, 10 mM Tris, pH 7.5, 1.5 mM MgCl2) and 1 unit of Taq polymerase (Pharmacia Biotech, Germany) supplemented with 0.01% gelatin, 0.2 mM of each dNTPs (Pharmacia Biotech, Germany), 50 ρmol of ISSR primers (Table 2), and 50 ng of total genomic DNA. Amplification was performed in a thermal cycler (Thermolyne Amplitron) programmed for 1 cycle of 2 min at 94°C; and 35 cycles of 30 secs at 94°C, 45 secs at 44°C, and 1.3 min at 72°C; followed by 20 min at 72°C.

**Table 2:** ISSR primers used in this study and a summary of the ISSR markers from *B. tabaci* samples

| Primer | Primer sequence | Amplified products | Fraction polymorphic fragments<sup>a</sup> |
| --- | --- | --- | --- |
| P02 | (ATCG)<sub>4</sub> | 0 | 0/0 |
| D12 | (GA)<sub>6</sub>CG | 5 | 1/5 |
| D14 | (CAC)<sub>3</sub>GC | 5 | 0/5 |
| D24 | (CA)<sub>6</sub>CG | 3 | 0/3 |
| HB13 | (GAG)<sub>3</sub>GC | 4 | 0/4 |
| HB14 | (CTC)<sub>3</sub>GC | 5 | 0/5 |
| UBC807 | (AG)<sub>8</sub>T | 6 | 5/6 |
| UBC810 | (GA)<sub>8</sub>T | 6 | 6/6 |
| UBC811 | (GA)<sub>8</sub>C | 6 | 4/6 |
| UBC823 | (AC)<sub>8</sub>C | 8 | 7/8 |
| UBC825 | (AC)<sub>8</sub>T | 8 | 7/8 |
| UBC826 | (TC)<sub>8</sub>A | 7 | 7/7 |
| UBC827 | (AC)<sub>8</sub>G | 12 | 12/12 |

<sup>a</sup> Number of polymorphic fragments/number of fragments amplified.

After completion of PCR, samples were cooled immediately to 10°C and stored at 4 °C until gel separation. A gel-loading solution (5 µl) was added, and 10 µl of the total production volume was resolved in 1.5% agarose in 1X TAE buffer for 1 h with a 100-bp ladder (Pharmacia, Germany) as the size standard. Gels were stained in ethidium bromide and images were recorded.

### ISSR-PCR Data analysis

Data were statistically analyzed by using a randomized complete block design with four replicates according to (27). The two growing seasons were analyzed separately. The least significant differences (LSD) test was used to compare means at the 5% level. Only differences significant at P≤0.05 are considered in the text.

Data of ISSR analysis were scored for computer analysis based on the presence or absence of the amplified products for each ISSR primer. If a product was present in a pest, it was designated “1”, if absent it was designated “0”. Pair-wise comparisons of *B. tabaci*, based on the presence or absence of unique and shared polymorphic products, were used to generate similarity coefficients based on the SIMQUAL module. The similarity coefficients were then used to construct a dendrogram by UPGMA (Un-weighted Pair-Group Method with Arithmetical Averages) using NTSYS-PC software version 2.0 (Exeter Software, New York) (28).

### Mitochondrial DNA amplification and Sequences

For mitochondrial cytochrome oxidase subunit I (mtCOI) sequences, DNA was extracted from nine individuals of *B. tabaci* and subjected to PCR amplification with the primer set according to (29) that include C1-J-2195 (5^/^ - TTGATTTTTTGGTCATCCAGAAGT-3^/^) and L2-N-3014 (5^/^– TCCAATGCACTAATCTGCCATATTA-3^/^). PCR products were sequenced (Judith Brown Lab., Univ. of Arizona, Tucson, USA). Other mtCOI sequences were obtained from NCBI (the public sequence databases).

### Sequence alignment and phylogenetic analysis

Using the CLUSTALW-based alignment tool, multiple sequence alignments were constructed by the available tool in MEGA version 5 (30). Produced sequences plus other close reference sequences obtained from the GenBank database for species from *B. tabaci* were loaded into Mega software for sequence alignments. The available nine sequences produced from this study were used to infer the relationship between them in comparison with different population samples from the public database and a phylogenetic tree was conducted with an initial approximate maximum likelihood.

### Statistical analyses

Obtained data were subject to statistical analysis using one-way analysis of variance (ANOVA) and means were compared using the Waller-Duncan post hoc multiple comparison test (31).

## Results

### Population density of *B. tabaci* immature colonized ten plant hosts

*Bemisia tabaci* population density varied according to the plant host type (Table 3). Data showed that cabbage (37.0±6.51, 24.2±6.8, and 23.0±3.08 eggs/cm2 at Al-Bukariyah, Ounen Al-Gawa, and Al-Midhnab, respectively) and cauliflower (34.2±6.23 eggs/cm2 at Al-Bukariyah) were the most favorable plants that helped the insect to colonize and deposited the highest number of eggs/cm2 than other hosts. Eggs number was also varied on the same plant host that cultivated at different locations. Cucumber came next with 27.6±1.84 eggs/ cm2. In contrast, lettuce was not a suitable host for insect oviposition which received 3.6±2.41 eggs/ cm2 at Agric. Exp. St., Faculty of Agriculture & Medicine Veterinary. Statistically, number of eggs varied significantly among plant hosts (F value = 16.66 with 9/40 df; Prob>F = <0.001) (Tab. 3).

**Table (3):** Host plants of *B. tabaci*, immature population density, and squash silverleafing symptoms at different Qassim.areas.

| Code | Location | Host plant | SSL Symptoms | *B. tabaci* population density/cm<sup>2</sup> |  |
| --- | --- | --- | --- | --- | --- |
|  |  |  |  | Eggs | Nymphs |
| V1 | Agri. Exp. St., Qassim Univ. | Lettuce | + | 3.6±2.41 g | 4.8±1.48 f |
| V4 | Agri. Exp. St., Qassim Univ. | Cucumber | ++ | 27.6±1.84 bc | 14.6±3.78 b |
| V7 | Ouen Al-Gawa | Cabbage | + | 24.2±6.8 cd | 13.2±1.48 bc |
| V6 | Ouen Al-Gawa | Potato | + | 16.8±3.65 def | 10.6±2.07 c |
| V10 | Al-Melada | Tomato | + | 8.4±3.46 g | 5.4±2.07 f |
| V9 | Unayzah | Eggplant | + | 16.3±3.96 ef | 9.4±3.07 cd |
| V2 | Al-Bukariyah | Cabbage | ++ | 37.0±6.51 a | 18.8±3.27 a |
| V3 | Al-Bukariyah | Cauliflower | + | 34.2±6.53 ab | 22.2±6.14 a |
| V5 | Al-Badai | Sweet pepper | + | 9.4±2.70 fg | 6.4±2.30 df |
| V8 | Al-Midhnab | Cabbage | ++ | 23.0±3.08 cde | 10.6±2.30 c |
| **Statistical Analysis** |  |  |  |  |  |
| F value |  |  |  | 16.66 with 9/40 df | 17.13 with 9/40 df |
| Prob> F |  |  |  | <0.0001 | <0.0001 |
| LSD at 5% |  |  |  | 7.452 | 3.993 |
| LSD at 1% |  |  |  | 10.036 | 5.378 |

• (-) No SSL symptoms were observed.
Means within a column followed by the same letter are not significantly different at the 5% level. ANOVA P < 0.0001 followed by Student-Newman-Keuls, P < 0.05.

Regarding population density of *B. tabaci nymphs*, cauliflower at Al-Bukariayah harbored the highest (22.2±6.14 nymphs/cm2), followed by cabbage at Al-Bukariayah (18.8±3.27 nymphs/cm2) and cucumber at Agric. Exp. St., Faculty of Agriculture & Medicine Veterinary (14.6±3.78 nymphs/cm2). On the other hand, lettuce was not a suitable host for insect oviposition that received 4.8±1.48 eggs/ cm2 at Agric. Exp. St., Faculty of Agriculture & Medicine Veterinary. Statistically, number of eggs varied significantly among plant hosts (F value = 17.13 with 9/40 df; Prob>F = <0.001) (Tab. 3).

#### I. Squash silverleaf (SSL) Symptoms

Data revealed that *B. tabaci* populations collected from the different agricultural areas at Qassim region have an ability to induce the squash silvering leaf symptoms (Table 3). Some of *B. tabaci* populations caused slight symptoms of SSL, with varying degrees of SSL symptoms. Squash silverleafing symptom (SSL) is a sign of kind of plant physiological disorders resulting as feeding activity of *B. tabaci* biotype “B” only. Therefore, it could be concluded that some of *B. tabaci* populations that invade Qassim agricultural areas are “B” biotype and which others may be indicated to other biotypes.

#### II. Cross mating

Experimental trials were carried to study the potential interbreeding among 10 populations of Collected *B. tabaci* from 8 areas at Qassim, to determine their compatible genetic structure. Table (4) summarized data of cross mating among the populations. Data revealed that cross mating has occurred among all populations, but few of them only produced both sex females and males while others were produced the only male. The groups that have higher similarity in genetic structures were able to lay fertilized and unfertilized eggs, which produce both females and males. Moreover, interbreeding among *B. tabaci* populations, that had a similar genetic structure or in a close related relationship, produced huge numbers of females than others (Table 4).

**Table (4):** Average number of laid eggs and sex ratio as a result of cross mating between 10 populations of *B. tabaci* collected from different plant hosts and areas at Qassim region.

| **Biotypes** | **V1** |  | **V2** |  | **V3** |  | **V4** |  | **V5** |  | **V6** |  | **V7** |  | **V8** |  | **V9** |  | **V10** |  |
| --- | --- | --- | --- | --- | --- | --- | --- | --- | --- | --- | --- | --- | --- | --- | --- | --- | --- | --- | --- | --- |
|  | Eggs | Sex | Eggs | Sex | Eggs | Sex | Eggs | Sex | Eggs | Sex | Eggs | Sex | Eggs | Sex | Eggs | Sex | Eggs | Sex | Eggs | Sex |
| **V1** | 210<br>±10.2 | ♂♀<br>1:2 |  | ♂ |  | ♂ | 144.1<br>±6.2 | ♂♀<br>1:2 |  | ♂ | 115.7±<br>11.3 | ♂♀<br>1:1.5 | 95.2±<br>9.6 | ♂♀<br>1:1.5 |  | ♂ | 105±<br>12.4 | ♂♀<br>1:1.7 | 119±<br>10.8 | ♂♀<br>1:1.8 |
| **V2** |  | ♂ | 221.2±<br>12.7 | ♂♀<br>1:2 | 112.1±<br>7.6 | ♂♀<br>1:1.5 |  | ♂ |  | ♂ |  | ♂ |  | ♂ |  | ♂ |  | ♂ |  | ♂ |
| **V3** |  | ♂ | 112.1±<br>7.6 | ♂♀<br>1:1.5 | 225±<br>1.9 | ♂♀<br>1:2 |  | ♂ |  | ♂ |  | ♂ |  | ♂ |  | ♂ |  | ♂ |  | ♂ |
| **V4** | 144.1<br>±6.2 | ♂♀<br>1:2 |  | ♂ |  | ♂ | 199.1±<br>11.5 | ♂♀<br>1:1.5 |  | ♂ |  | ♂ |  | ♂ |  | ♂ | 95.1±<br>3.1 | ♂♀<br>1:1.5 | 77.3±<br>4.2 | ♂♀<br>1:1.5 |
| **V5** |  | ♂ |  | ♂ |  | ♂ |  | ♂ | 175±<br>9.5 | ♂♀<br>1:2 |  | ♂ |  | ♂ |  | ♂ |  | ♂ |  | ♂ |
| **V6** | 76.2±<br>1.3 | ♂♀<br>1:1.5 |  | ♂ |  | ♂ | 87.2±<br>4.1 | ♂♀<br>1:1.5 |  | ♂ | 189.4±<br>10.7 | ♂♀<br>1:2 | 96.2<br>±3.2 | ♂♀<br>1:1.2 |  | ♂ | 94.6<br>±4.2 | ♂♀<br>1:1.4 | 84.7<br>±5.2 | ♂♀<br>1:1.6 |
| **V7** | 95.2±<br>9.6 | ♂♀<br>1:1.5 |  | ♂ |  | ♂ | 66.2±<br>2.2 | ♂♀<br>1:1.5 |  | ♂ | 96.2<br>±3.2 | ♂♀<br>1:1.2 | 201±<br>8.8 | ♂♀<br>1:2 |  | ♂ | 96.3<br>±4.2 | ♂♀<br>1:1.4 | 81.2<br>±3.8 | ♂♀<br>1:1.5 |
| **V8** |  | ♂ |  | ♂ |  | ♂ |  | ♂ |  | ♂ |  | ♂ |  | ♂ | 197±<br>10.1 | ♂♀<br>1:2 |  | ♂ |  | ♂ |
| **V9** | 105±<br>12.4 | ♂♀<br>1:1.7 |  | ♂ |  | ♂ | 95.1±<br>3.1 | ♂♀<br>1:1.5 |  | ♂ | 94.6<br>±4.2 | ♂♀<br>1:1.4 | 96.3<br>±4.2 | ♂♀<br>1:1.4 |  | ♂ | 210±<br>11.4 | ♂♀<br>1:2 | 73.3<br>±5.9 | ♂♀<br>1:1.5 |
| **V10** | 119±<br>10.8 | ♂♀<br>1:1.8 |  | ♂ |  | ♂ | 77.3±<br>4.2 | ♂♀<br>1:1.5 |  | ♂ | 84.7<br>±5.2 | ♂♀<br>1:1.6 | 81.2<br>±3.8 | ♂♀<br>1:1.5 |  | ♂ | 73.3<br>±5.9 | ♂♀<br>1:1.5 | 222.3±<br>10.9 | ♂♀<br>1:2 |

\* Average No. of *B. tabaci* eggs shown in this table is the fertilized eggs only (as a result of cross mating between biotypes which produced females and males, as well as, the symbol of sex ratio).

The importance of interbreeding among *B. tabaci* population is a clear evidance that individuals with cytoplasmic compatibility and belonging to the same genotype produce more females than males, and vice versa, The important results of the cross mating, by females paired with males of different genetic structures, were reduced population, a notable increase in male progeny proportion, fewer eggs produced, and no difference in egg number/unmated females and females paired with males of the similar gene structure.

#### III. ISSR-PCR analysis

DNA analysis using ISSR-PCR of 10 *B. tabaci* populations, as well as biotype “B” as a control, collected from different hosts and locations at Qassim region indicated that these insects were genetically varied. In comparison with *B. tabaci* biotype B as a control, data demonstrated that the genetic structures of selected insects found at the Qassim region have differed from biotype B. This means that *B. tabaci* which spread and infested Qassim vegetables under greenhouses varied genetically from the most famous biotype called “B”. Obtained results may be lead to scientific evidence on the presence of new biological biotypes from this insect that differed completely from the biotype “B”. Additionally, based on ISSR-PCR analysis, *B. tabaci* populations (10 populations) collected from eight agricultural areas in Qassim region could be divided into 4 groups based on ISSR analysis (Fig 1) (Table 5). The genetic similarity of biotype “B” was varied by about 52% from Qassim population of *B. tabaci* (Fig. 2). Similarity among the collected *B. tabaci* population from Qassim region was about 68% but varied among the four groups according to location and hosts. Bemisia populations in group 1 which included 6 populations (V1 & V4, lettuce, and cucumber from Agric. Exp. St.; V6 & V7, potato and cabbage from Ouen-Al-Gwa’a; V9, eggplant from Uniyzia and V10, tomato from Melada) were similar by 80%. Group 2 which included 2 populations (V2, cabbage, and V3, cauliflower from Al-Bukariyah were similar by 84%. Group 3 that included one population collected from cabbage from Al-Midhnab was similar to group 1 and 2 in average72%, while group 4 included one population infected pepper from Al-Badai was similar to the other groups by 68% (Fig. 2). Scientifically, the four genetic groups are not identical to the B biotype, and maybe new genetic structures that not known before at Qassim in particular and KSA in general.

**Table 5.**
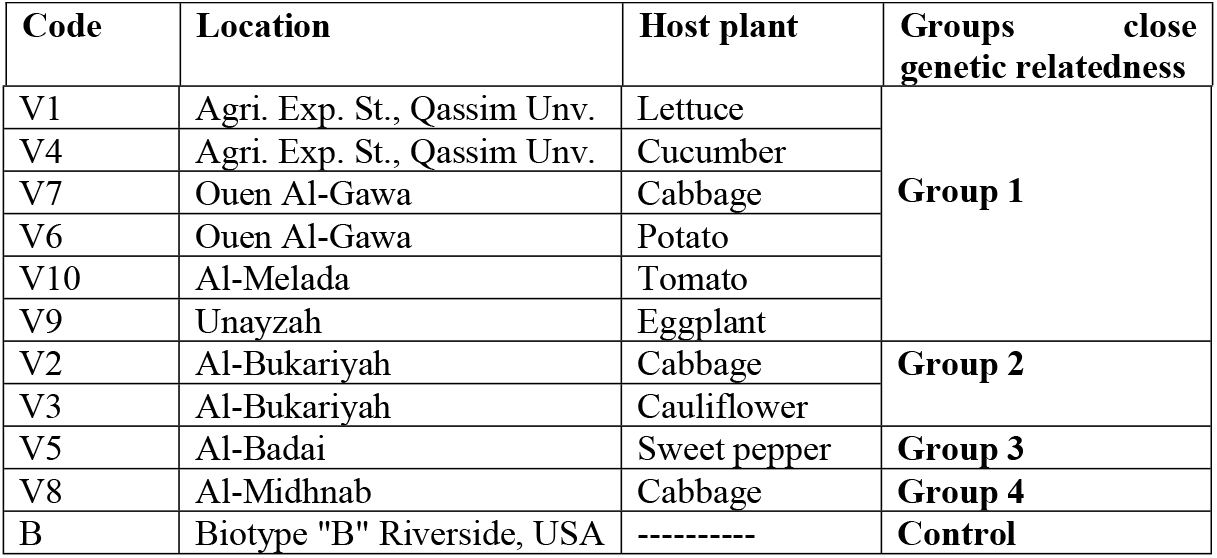
Sample code, location, plant host, and group classification depending on ISSR are shown as following.

**Fig 1.**
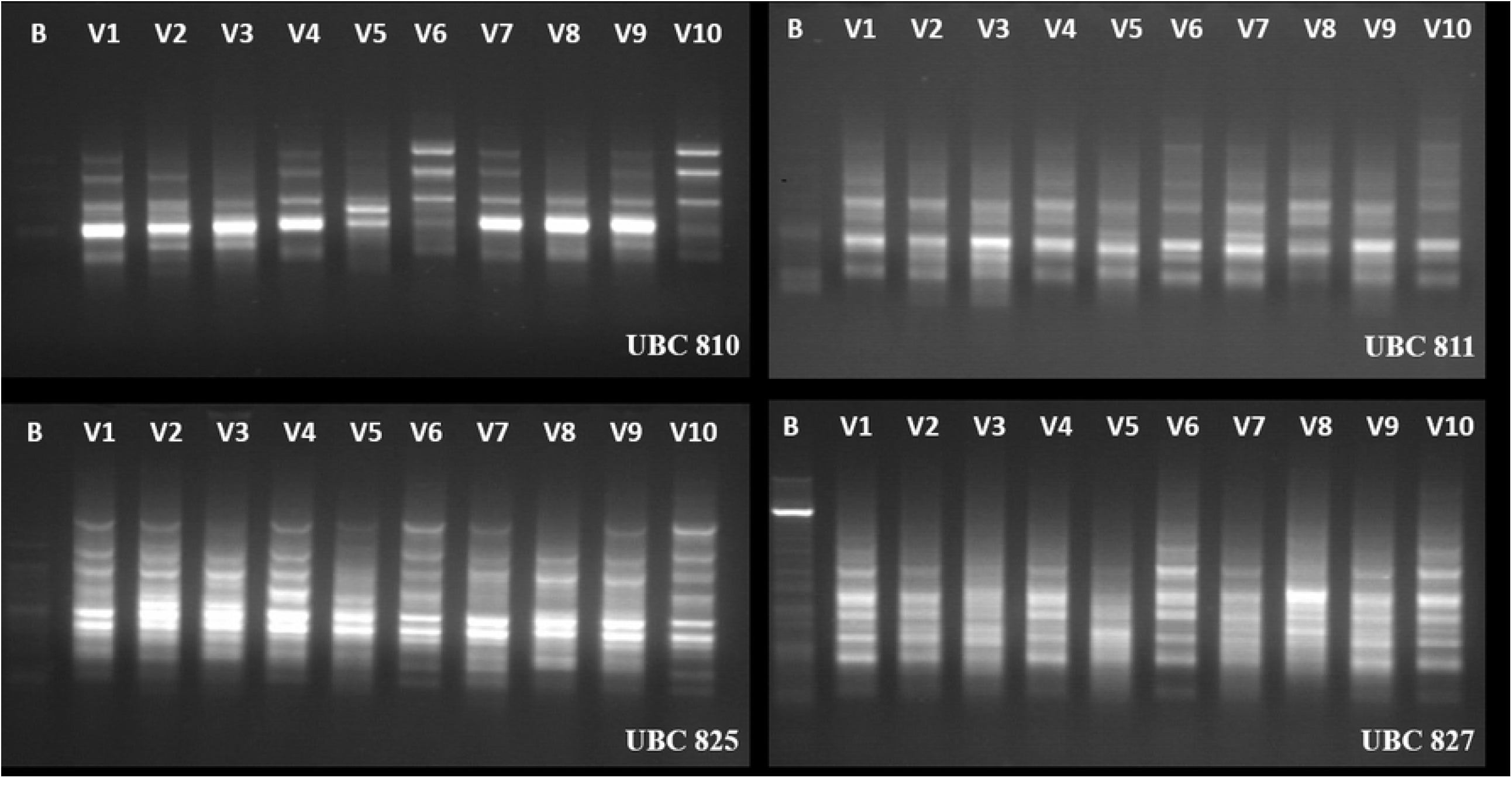
ISSR-PCR results of 11 DNA samples of *B. tabaci*, amplified by primers UBC 810, UBC 811, UBC 825, and UBC 827.

**Fig 2.**
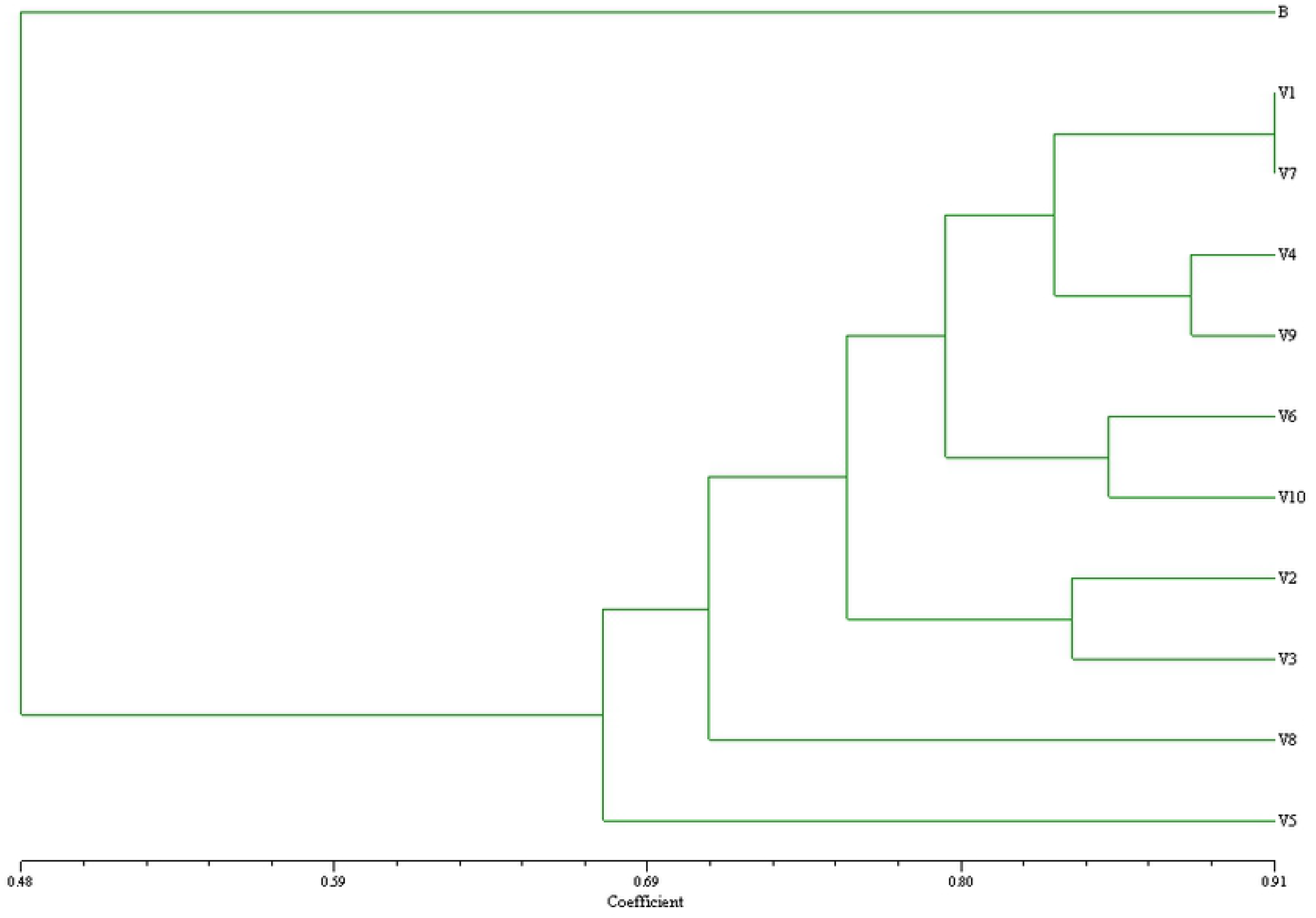
Phylogenetic analysis of ten *B. tabaci* populations ISSR-PCR sequences at Qassim Agricultural region. Sample code, location, and plant host shown as following.

### Phylogenetic analysis of *Bemisia tabaci* by Mitochondrial COI (mtCOI)

A total of twenty-two whitefly sequences from *B. tabaci*, were collected from seven locations and analyzed in our study (table 1). Phylogenetic analyses revealed two major putative groups in addition to out-group sequences from *B. tabaci* (*i*.*e*. EegTB12 (8-3); B biotype sequences from Spain, Uganda; *T. ricini* Egypt 2; Ivory coast cassava) (Fig 3). Otherwise, the first group involve two clades, the first clade had Asia I, II (biotypes from India and Pakistan) whereas the other clade contains biotypes from the Americas (i.e. Arizona, California, and biotype A). The second group belongs to North Africa, the Mediterranean, and the Middle East region. This group comprised two haplotypes divided into two clades belong to B and Q biotypes. The Q biotype has species from Turkey, Morocco, Israel, and Ivory Coast Okra, these individuals were found to be closely related to B biotypes. On the other hand, B biotype clade includes almost all submitted sequences in this study. Approximately, twenty-one sequences were located and clustered with B biotype clade (*i*.*e*. EegTb12 (Eggplant), EsqTb12 (Squash), EcabTb12 (Cabbage), EcauTb12 (Cauliflower), EconTb12 (Convolvulus), EtoTb12 (Tomato)) whereas EegTB12 (8-3) exhibited the highest diversity and located in the out-group species.

**Fig 3.**
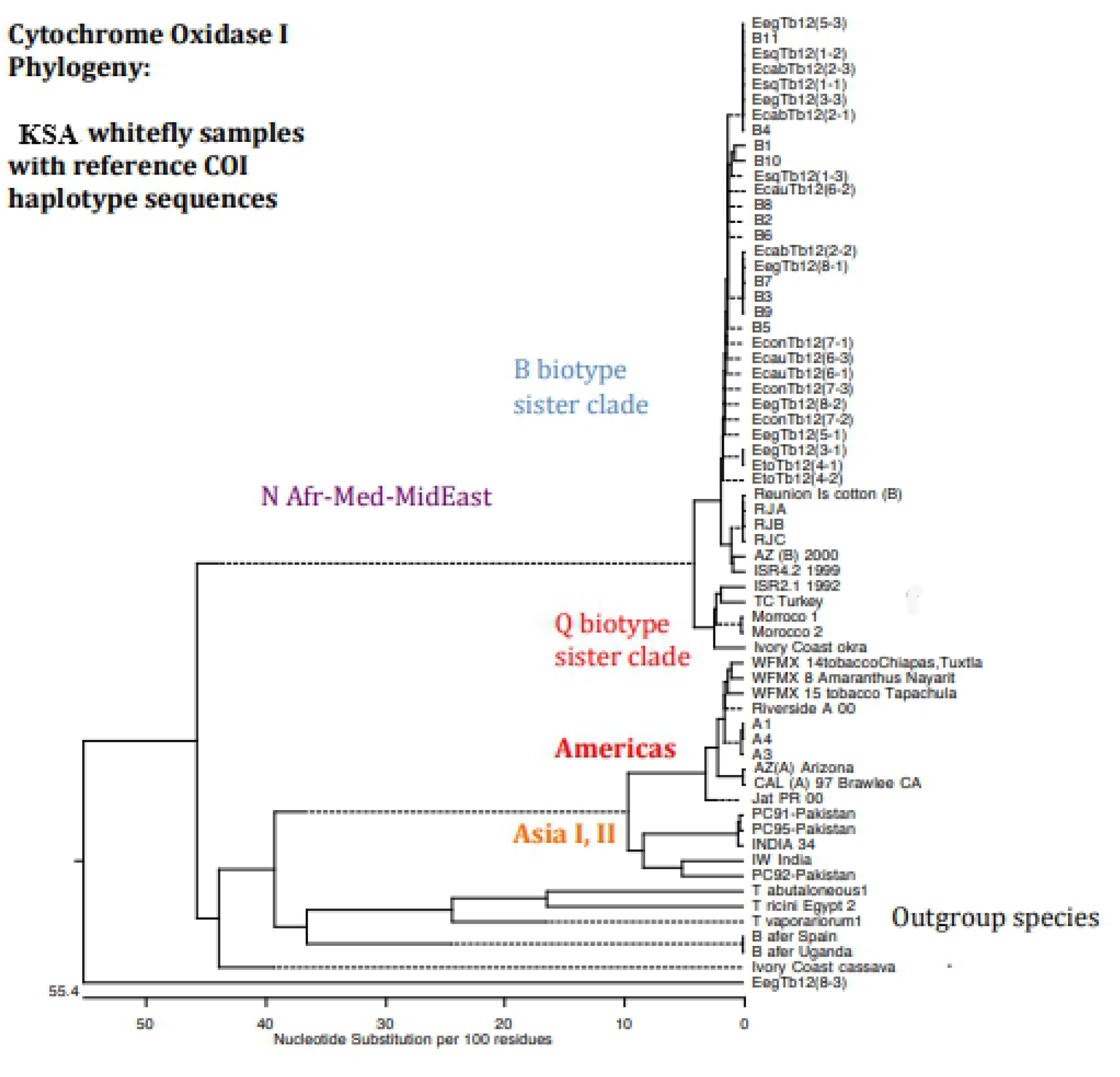
Phylogenetic tree indicating the relationships between twenty-two mitochondrial COI DNA sequences of *B. tabaci* in addition to sequences obtained from the GenBank database. The tree was constructed by an approximate maximum likelihood phylogeny.

## Discussion

*Bemisia tabaci* is a dramatic example of insects that have remarkable variations among its populations as a result influence of many factors which affect alone or in combination with others such as; host adaptations, weather factors, geographic areas, and insecticides (12) Variation of *B. tabaci* genetic structure patterns within and among host plants and geographic regions were discussed carefully in the last 30 years (32)(16)(8). Accordingly, *B. tabaci* is a species complex, which its status has been escalated over the past 30 years and became one of the worst 100 pests in the world. Until now, at least 24 biotypes belongs to 7 genetic groups of the pest were determined (15).

In the current study, population densities of *B. tabaci* (eggs and nymphs) collected from 10 plant hosts at the Qassim region (Table 3) were significantly varied according to its host plant and among agricultural areas. Because all agricultural areas are located in the Qassim region and have the same weather condition, so, the location had a slight influence on the population densities. Therefore, the variations among its populations may be due to the effect of host type, and this is visible and apparent among the same location, which had different plant hosts (Table2). These results go hand in hand with (33)(34)(35). Finally, there are over twenty Bemisia biotypes distributed worldwide (13). Biotypes distribution and their plant hosts varied from one country to another. The same country and the same plant hosts may be infested with more than one biotype

The most visible sign of the newly virulent *B. tabaci* biotype “B” is the unusual discoloration or “silvering” of squash leaves. (36)(37)(3)(33) used SSL symptoms as an indicator of the presence of *B. tabaci* biotype “B” on the Egyptian plant hosts. The results differentiated between two different genetic biotypes namely, biotype “B” and biotype “Q”.

Data obtained in this study declared that all *B. tabaci* populations at Qassim haven’t the ability to induce the squash silverleafing symptoms (SSL) or induce very slight symptoms (Table 3). Therefore, it could declare that *B. tabaci* biotype “B” had no populations at Qassim regions, and populations that invade and infest horticulture plants at Qassim may be another biotype or biotypes. This toxicogenic disease was linked to plant disorders as a result of biotype “B” feeding (4)(19). B-type nymphs are responsible for inducing the silverleaf symptoms (38). The silverleaf disorder is of economic importance because the affected tissue reduces chlorophyll and high light reflectance that may result in lower yield (39).

The sweet potato whitefly, *B. tabaci* is haplodiploid. Fertilized eggs give rise to diploid females, whereas, males are produced from unfertilized eggs. In a diploid (2N), females are produced from fertilized eggs, while haploid (1N) produce only males from unfertilized eggs (40). Moreover, when males’ number in a population decline, fertilization is reduced, leading to an increased number of male offspring in the next generation. As a result, sex ratio bias is frequently reported for alternate generations of this whitefly (41)(42). This is in agreement with the obtained data (Table 4) when the interbreeding was carried out among the whitefly populations collected from Qassim horticulture plants. Populations that have the same genetic structure or in a high degree of similarity, were produced females and males, with higher sex ratios in favor of females. On the Contrary, populations that have no similarities in their genetic structures gave high numbers of males and no females were detected (Table 4).

(43) reported three explanation for this case: 1) the distracting male hypothesis in which mating pairs made up of different biotypes spent more time to courtship and less time to egg-laying than single biotype pairs; 2) the single-locus complementary sex determination model in which the production of non-viable diploid male zygotes may explain the reduction in eggs laid and; 3) cytoplasmic incompatibility between biotypes caused by Wolbachia. Finally, the results also suggest the geographical distribution of related biotype clusters both overseas and in Australia may be explained by between-biotype interactions leading to the formation of parapatric populations. Many studies declared it is not possible that the interbreeding between different biotypes of *B. tabaci* or varied in their same genetic structures or if happen it will produce males only. In this respect, (44) by mating experimental was able to differentiate among six cryptic species in the *B. tabaci* complex

The results also pointed out that ISSR-PCR analysis might be useful in distinguishing closely related organisms particularly biotypes. Figures 1 demonstrated that populations of *B. tabaci* in Qassim region hadn’t the same genetic structures. ISSR-PCR analysis represents different genetic structures and this could be gathered by phylogenetic relationships into 4 genetic groups. These groups were similar to “B” biotype with 48% while the similarity degree among others ranged from 67-94%. This observation suggests that it is most aptly considered a species complex. The complex comprises multiple genetic haplotypes and many well-characterized, behaviorally distinct variants, referred to as biotypes (12)(8).

Regarding Cytochrome oxidase subunit I analysis, our results preview that all the sequenced individuals were clustered in one clade (B biotype) except one sequence (EegTB12 (8-3)) which was the most diverts and had been considered as out-group. In North Africa and the Middle East, both Q and B type are widespread while in America, the A-type is dominant. (45) Characterized *B. tabaci* populations occurring on cassava in the Central African Republic. The authors detected a high level of genetic diversity. Molecular studies of *B. tabaci* provided significant evidence for polymorphism degree between populations in selected geographic regions from CAR. In another study, 59 *B. tabaci* populations were detected to belong to either biotype B or Q concerning PCR-RFLP patterns of the mtCOI gene for the purpose to determine a diagnostic microsatellite marker. Two restriction patterns were revealed (414, 162, 144 bp) and (631,144 bp) and found to be related to biotype B and Q respectively (46). In Turkey, populations and various biotypes of the cotton whitefly (*B*. tabaci) were collected and the genetic diversity was determined using mitochondrial cytochrome oxidase subunit 1 (mtCOI) sequences. Analysis of sequences by phylogenetic revealed that *B. tabaci* populations of Turkey were divided into two groups clustering around the main Mediterranean groups, (B and Q biotypes) and Middle East-Asia Minor 1 (47). (48) in South Sudan, studied the genetic variability of *B. tabac*i that infect cassava and sweet potato.

In conclusion, squash silverleaf will become a perfect tool of *Bemisia* classification when combined with ISSR-PCR analysis and interbreeding between biotypes. The authors emphasized that the cross-mating, genetic analysis, morphological and physiological examination must be done first to identify Bemisia biotypes. Therefore, this study declares that *B. tabaci* invade and infest Qassim horticulture haven’t the same genetic structures and not A or B biotypes according to ISSR technique but belong to B biotype when Cytochrome oxidase subunit I was used.

